# Selective neuronal vulnerability in Alzheimer’s disease: a network-based analysis

**DOI:** 10.1101/499897

**Authors:** Jean-Pierre Roussarie, Victoria Yao, Zakary Plautz, Shirin Kasturia, Christian Albornoz, Eric F Schmidt, Lars Brichta, Alona Barnea-Cramer, Nathaniel Heintz, Patrick Hof, Myriam Heiman, Marc Flajolet, Olga Troyanskaya, Paul Greengard

## Abstract

A major obstacle to treating Alzheimer’s disease (AD) is our lack of understanding of the molecular mechanisms underlying selective neuronal vulnerability, which is a key characteristic of the disease. Here we present a framework to integrate high-quality neuron-type specific molecular profiles across the lifetime of the healthy mouse, which we generated using bacTRAP, with *postmortem* human functional genomics and quantitative genetics data. We demonstrate human-mouse conservation of cellular taxonomy at the molecular level for AD vulnerable and resistant neurons, identify specific genes and pathways associated with AD pathology, and pinpoint a specific functional gene module underlying selective vulnerability, enriched in processes associated with axonal remodeling, and affected by both amyloid accumulation and aging. Overall, our study provides a molecular framework for understanding the complex interplay between Aβ, aging, and neurodegeneration within the most vulnerable neurons in AD.

## Introduction

Selective neuronal vulnerability is a shared property of most neurodegenerative diseases^1^. The molecular basis for this selectivity remains unknown. In the early stages of Alzheimer’s Disease (AD), the most common form of dementia, clinical symptoms (such as memory loss) are caused by the selective degeneration of principal neurons of the entorhinal cortex layer II (ECII), followed by CA1 pyramidal cells in the hippocampus. In contrast, other brain regions, such as the primary sensory cortices, are relatively resistant to degeneration until later stages of the disease^2-8^.

AD is characterized by two major pathological hallmarks: accumulation of the Aβ peptide (the main constituent of amyloid plaques) and formation of neurofibrillary tangles (NFT, aggregates of hyperphosphorylated tau proteins which are thought to occur downstream of Aβ accumulation). Amyloid plaques do not accumulate in discrete brain areas. Rather, they are relatively widespread across most regions of the neocortex, followed by the entorhinal cortex and hippocampus of AD patients^9,10^. In contrast, NFTs exhibit the same regional pattern as neurodegeneration^11-13^. The co-occurrence of NFTs and neurodegeneration, as well as the fact that the best pathological correlate for clinical symptoms to date is the extent of NFT formation^14-16^, highlight the importance of tau pathology. Genetic analyses have revealed the importance of microglia in the disease. Yet the molecular drivers for the neuronal component of the pathological cascade that leads from Aβ accumulation to NFT formation and neurodegeneration are still largely unknown. While there might be regional differences in microglia identity, recent evidence suggest that microglia regional particularities are mainly driven by regional differences in the neighboring neurons^17^.

To understand and model cell type-specific vulnerability in AD, we must thus gain insight into the molecular-level differences between healthy vulnerable and resistant neurons that predispose some neurons, before any pathological process becomes visible, to develop tau pathology much faster than others. This requires high quality cell type-specific profiles of both vulnerable and resistant neurons. While some neuron types of relevance to AD were profiled in a mouse hippocampal study^18^, the most vulnerable neuron type in early AD (ECII) has not previously been studied *ex vivo.* In humans, Small *et al*. profiled whole EC and dentate gyrus (DG) in control and AD patients^19^, and the Allen Brain Atlas (ABA) provides a large dataset for a number of human brain regions^20^, but neither of these studies are cell type specific. A comprehensive dataset of neuron-specific AD-relevant profiles has been generated by Liang *et al*^21^. While valuable, human samples, including those in the studies cited above, are inevitably subject to degradation and *postmortem* changes and, in the context of AD, do not allow for direct probing of the effect of aging and Aβ accumulation on gene expression.

Furthermore, a key challenge to achieving a molecular understanding of selective neuronal vulnerability in AD is that vulnerability and pathology are likely not simply the result of a few genes acting in isolation. Previous work examining whole brain lysates from AD patients and non-demented individuals ^22-24^ demonstrated the promise of network analyses in AD, but these studies were limited to larger brain regions and thus could not address cell type-specific vulnerability. Deciphering the pathological cascade requires cell type-specific systems-level analysis and modeling of the complex molecular interactions that underpin the vulnerability of specific neurons to AD. Here, we provide the first molecular framework to understand the interactions between age, Aβ, and tau within neurons. Our approach (Fig. 1a) integrates the precision of cell type-specific profiling across age in the non-diseased mouse with computational modeling of human neuronal-omics (e.g., expression, interaction) data. It also combines this network modeling with human disease information from quantitative genetics data, ensuring relevance for human AD biology. The neuron-specific networks are available for download and exploration in an interactive web interface (alz.princeton.edu).

**Figure 1:**
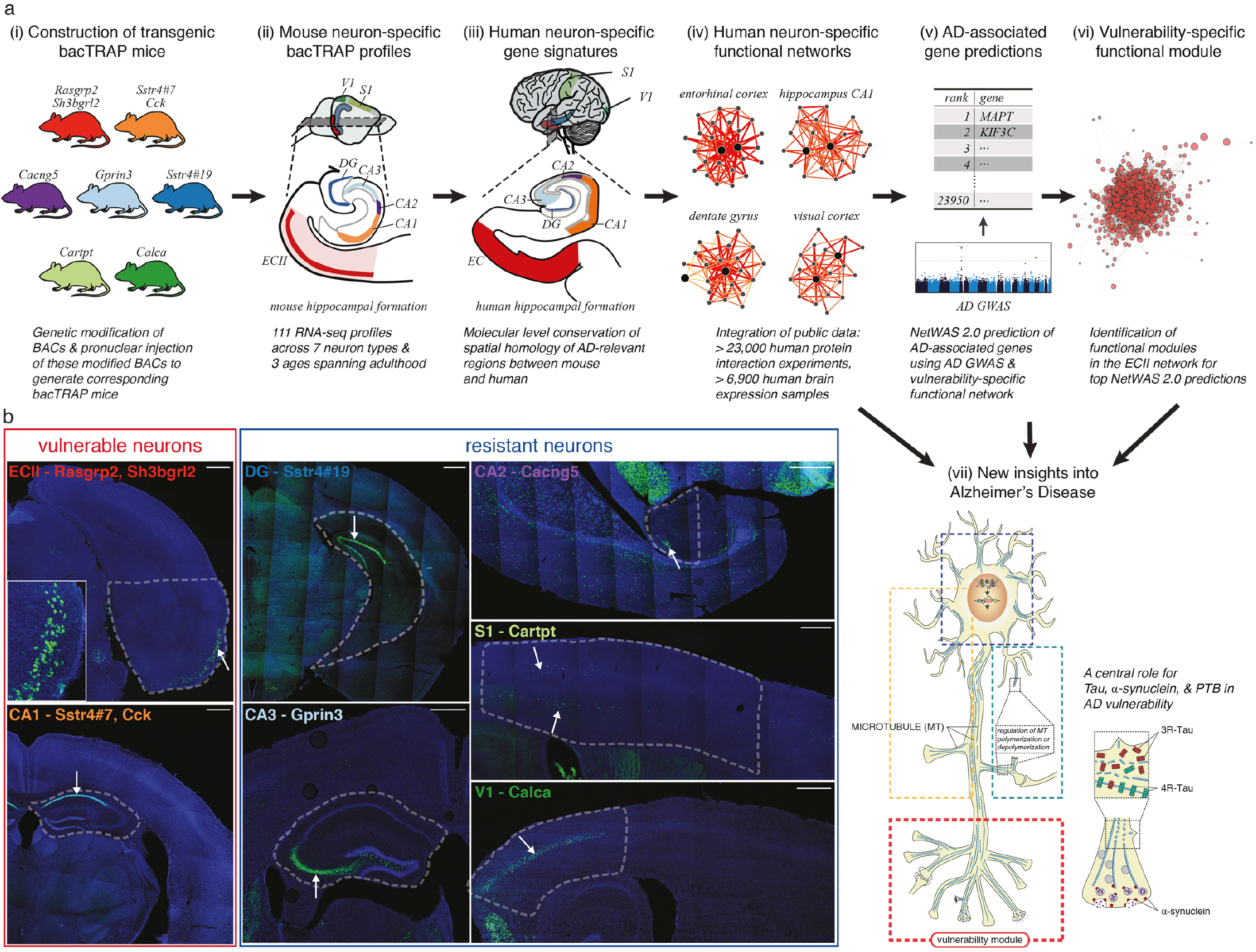
An integrative experimental genomics and bioinformatics framework, combining mouse and human data, to identify genes and pathways involved in Alzheimer’s Disease. **a. Overview of our framework.** (i) To obtain molecular profiles of 7 types of neurons most vulnerable and most resistant to AD using the bacTRAP technology, we constructed bacTRAP mice for each of them (see b). (ii) 111 neuron-specific high quality *ex vivo* expression profiles were obtained for each neuron type at three different ages (5 months, 12 months, 24 months), using the bacTRAP technology with these mice followed by deep sequencing: bacTRAP allows for fast isolation of actively translated RNA with only minimal alterations of mRNA content after the death of the animal, and quantitative assessment of gene expression over a large range of expression levels. (iii) Using these data, we generated neuron-specific molecular signatures in mouse and human, and created a spatial homology map between the two organisms. (iv) We used these neuron-specific signatures to construct 7 neuron-specific functional networks, through Bayesian integration of a compendium of over 30,000 human experiments. (v) We identified AD-associated genes by combining the network for the most vulnerable neuron (ECII) with an AD tau pathology GWAS study^42^ using our NetWAS 2.0 machine learning approach. (vi) These genes form distinct functional modules in the vulnerable neuron-specific network, with one module in particular capturing vulnerability-specific signal. (vii) Our analyses point to the involvement of neurotransmitter release and axonogenesis in AD vulnerability, as well as a central role for the regulation of tau and α-synuclein by the RNA-binding protein PTB. Overall, we map AD-associated processes and their potential regulation by aging and Aβ in ECII neurons, providing the first molecular dissection of the AD pathological cascade within vulnerable neurons. **b. bacTRAP transgenic mice generated for the molecular profiling of vulnerable and resistant neurons.** For each line, brain sections were stained with an anti-eGFP antibody (green) and counterstained with DAPI (blue). Genes whose regulatory regions we used for driving eGFP-L10a expression are indicated in each frame. For ECII and CA1, we show a section from the Rasgrp2- and from the Sstr4#7-bacTRAP line respectively, but also used Sh3bgrl2- and Cck-bacTRAP lines for subsequent analyses. The dashed line delineates the brain region dissected out for bacTRAP. Arrows point to the neurons of interest, which overexpress eGFP-L10a. Scale bar: 500 μm (ECII: principal neurons of the layer II of the entorhinal cortex, CA1: pyramidal neurons of hippocampus CA1, CA2: pyramidal neurons of hippocampus CA2, CA3: pyramidal neurons of hippocampus CA3, DG: granule neurons of dentate gyrus, S1: pyramidal neurons of primary somatosensory cortex, V1: pyramidal neurons of primary visual cortex)

## Results

### Cell type-specific profiling of mouse neurons with differential vulnerability to AD

To investigate the selective vulnerability of neurons in AD, we generated cell type-specific expression profiles spanning the entirety of adulthood for vulnerable and resistant neurons using the bacTRAP (Bacterial Artificial Chromosome – Translating Ribosome Affinity Purification) technology in normal mice^25,26^. The bacTRAP technology enabled us to assay AD-relevant neuronal cell types with genome-wide coverage, measure transcripts *ex vivo* (as opposed to *postmortem*), and specifically capture actively translated (rather than all transcribed) genes.

We focused on the vulnerable principal neurons of ECII and pyramidal neurons of CA1, and on five types of resistant neurons, namely pyramidal neurons of CA2, CA3, primary visual cortex (V1), primary somatosensory cortex (S1), and granule cells of DG. Specifically, we constructed different transgenic mouse lines for each type of neuron, overexpressing the ribosomal protein L10a fused to the green fluorescent protein (GFP) under the transcriptional control of a driver specific to that type of neuron (Fig. 1b, Supplementary Note). The bacTRAP procedure then consists of immunoprecipitation of GFP-tagged polysomes from GFP-L10a-expressing cells, thus isolating actively translated neuron-specific mRNAs for RNA-sequencing. Previous work using bacTRAP or similar technologies (e.g., RiboTag) has demonstrated the strong enrichment for cell-type specific signal from cells expressing the tagged ribosomal protein ^25-31^.

We first performed multidimensional scaling analysis of the resulting bacTRAP data and found that the samples (3-12 biological replicates per neuron type per age) clustered primarily by tissue location, as expected, with a clear separation between the ECII, hippocampal regions (CA1, CA2, CA3, DG), and neocortical regions (S1 and V1) (Fig. 2a). We further verified the expression patterns of known neuron-type specific markers (Fig. 2b) and identified the top enriched genes for each neuron type in our data (Fig. 2c, Supplementary Table 1). Comparisons with the semi-quantitative *in situ* hybridization (ISH) data in the ABA (Supplementary note, Supplementary Fig. 1) show that our data includes the cell type-specific signals in these datasets, while providing substantially higher regional and quantitative, genome-scale coverage. Thus, our approach provides a high quality, genome-wide assay of *ex vivo* neuron-type specific expression in AD-vulnerable and resistant regions of the brain.

**Figure 2:**
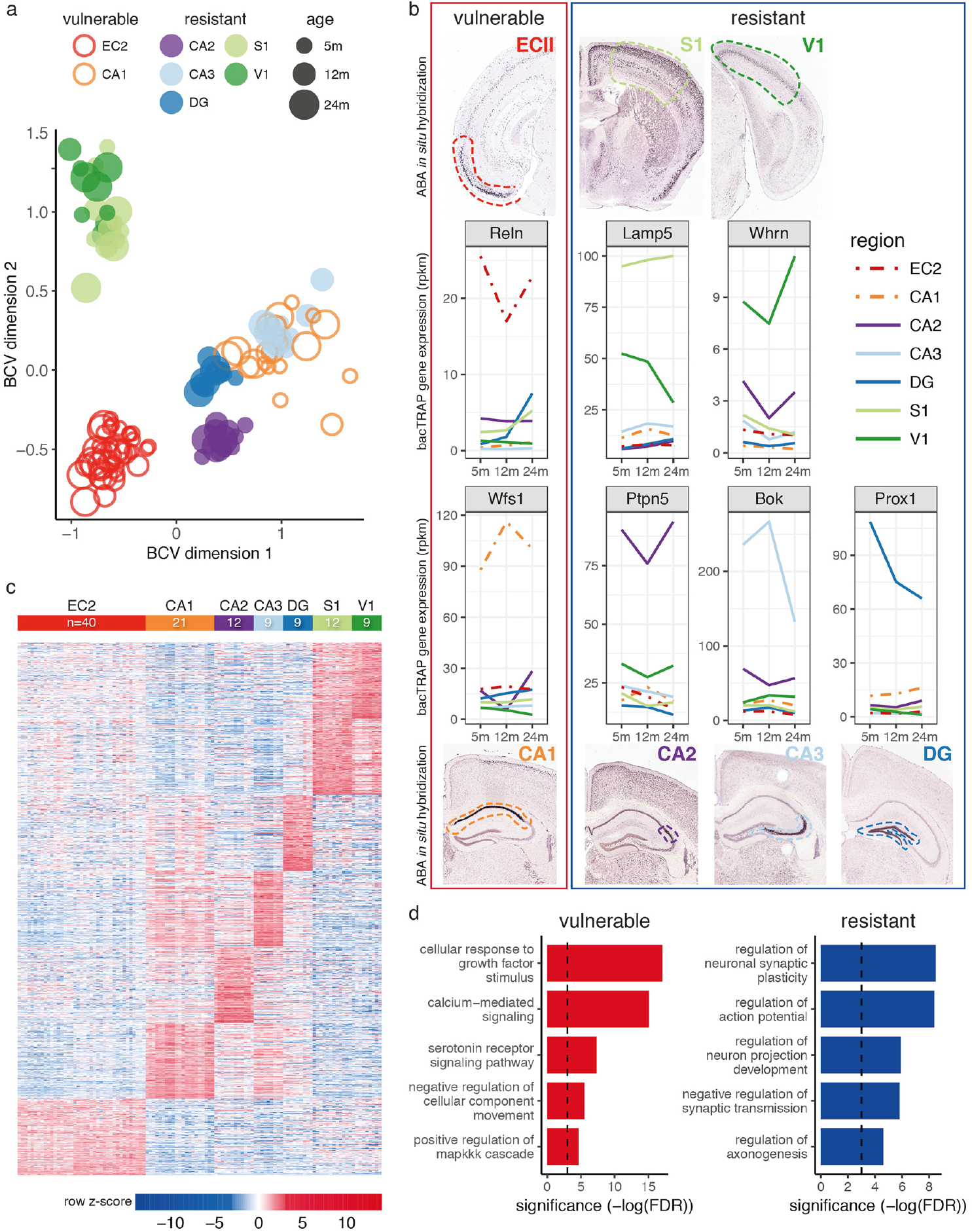
Molecular characterization of vulnerable and resistant neurons. **a. Multidimensional scaling analysis of all samples demonstrating clustering of samples by region of origin Each dot represents one sample (two mice pooled)**. Each color represents a different type of neuron. Red (ECII) and orange (CA1) dots correspond to AD vulnerable neurons. Purple (CA2), light blue (CA3), dark blue (DG), light green (S1), dark green (V1), dots correspond to resistant neurons. Increasing dot sizes represent increasing mouse ages (5 months, 12 months, 24 months). **b. Verification of our quantitative, neuron-specific RNAseq profiles for known markers by ISH (ABA)**. Expression of previously described cell type-specific markers across the seven types of neurons, Reln for ECII neurons^91^, Wfs1 for CA1 pyramidal neurons^92^, Ptpn5 (or Step) for CA2 pyramidal neurons^93^, Bok for CA3 pyramidal neurons^92^, Prox1 for dentate gyrus granule neurons^94^, Whrn for visual cortex layer IV^95^, and Lamp5 (or C20orf103) for somatosensory cortex layers II-III^31^. For each marker, we show the expression at 5 months, 12 months, and 24 months of age; each color represents a different type of neuron; we also show for each gene an ISH image from the ABA that shows expression in the corresponding neurons. Image credit: Allen Institute. **c. Heatmap of gene expression for the top 500 genes enriched in each neuron type**. For each gene (rows, grouped by neuron type in which they are enriched) and sample (columns, grouped by cell type, including all three different ages), the row-normalized log_2_(RPKM) is displayed, showing that hundreds of genes are enriched in each type of neuron. **d. Pathways enriched in vulnerable (red) and resistant (blue) neurons**, with their significance (-log(FDR)) of enrichment.

To characterize molecular signatures for AD vulnerable cells in the nondisease state, we compared gene expression profiles of ECII and CA1 neurons against the five AD-resistant neuron types in wild-type mice. Among the significantly enriched processes we found many AD-relevant pathways (Fig. 2d, Supplementary Table 2). Furthermore, known Alzheimer’s Disease genes were significantly enriched in vulnerable neurons (KEGG AD genes, Wilcoxon rank sum test, p-value < 9.41e-7). These results support the hypothesis that intrinsic properties of ECII and CA1 neurons, present even in healthy individuals, render these neurons as preferential substrates for the development of AD pathogenesis.

### Neuron-specific spatial homology between mouse and human

An important question for interpreting model organism studies of AD is whether the molecular identity of neurons is conserved between mouse and human. Previous comparisons using spatially resolved semi-quantitative ISH^32^ or transcriptomics and proteomics without cellular resolution^33,34^ have suggested that mouse and human regional expression patterns are correlated, but the conservation of expression across neuronal subtypes requires further exploration. In humans, fully quantitative data at cell type-specific resolution is lacking across the regions most relevant for AD. However, the discrete brain structure microarray data from the ABA^20^ captures enough regional specificity for an expression-based comparison between the seven mouse neuronal subtypes and 205 human brain regions. We calculated a spatial homology score between molecular signatures for each mouse neuron type and each human brain region, generating 1,435 pairwise spatial homology measurements. Remarkably, of all these possible mappings, we found a near perfect match between each mouse profile and its corresponding relevant human brain region (Fig. 3, Supplementary Table 3). This confirms the validity of leveraging the power of *ex vivo* neuron-specific molecular profiles in the mouse to gain relevant insight into the molecular characteristics of the most vulnerable neurons in human AD. While there are differences in lifespan and other factors relevant to AD that may facilitate the degeneration of human neurons^35^, our comparison supports the notion that physiological differences between vulnerable and resistant neurons are conserved. Our study provides, to our knowledge, the first systematic evidence that the molecular identity of AD-relevant cell types is conserved between the mouse and human brain. This supports our approach of combining the cell-type-specific signals in healthy mouse neurons with the AD-relevant signals in large collections of human data.

**Figure 3:**
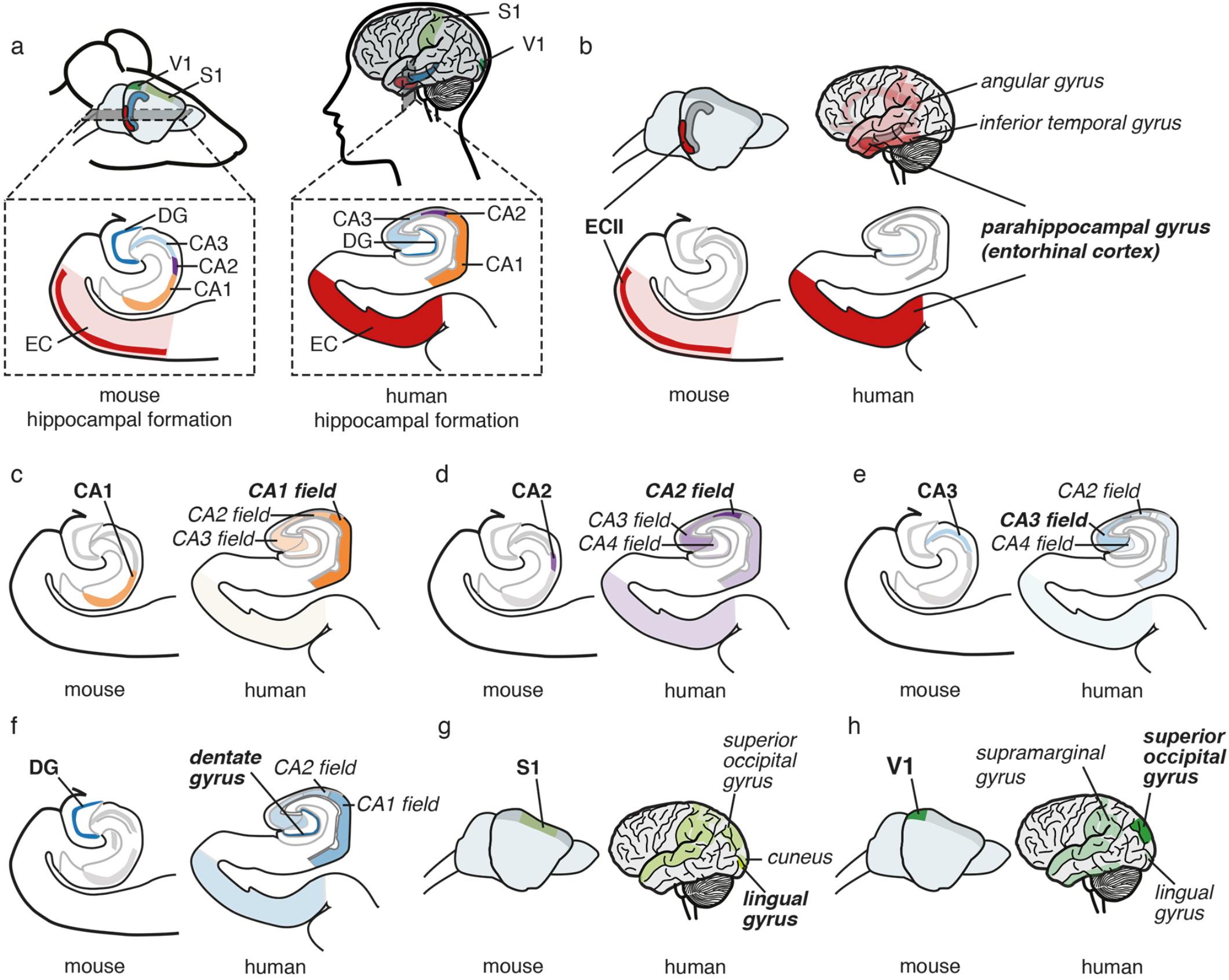
Conservation of molecular identity of seven AD-vulnerable and-resistant neuronal types between mouse and human. a. Location in the mouse and human brain of the seven brain regions included in this study (lateral view of the whole brain and close-up of the hippocampal formation). To validate the use of mouse profiles for the study of the human disease, we compared the molecular signatures of the mouse neurons derived from 111 mouse bacTRAP samples with 205 human brain region-specific expression profiles from the Allen Institute (b – h). For each mouse neuron type (b: ECII, c: CA1, d: CA2, e: CA3, f: DG, g: S1, h: V1), the human brain regions with molecular signatures closest to each mouse neuron type are highlighted (the more opaque the color of a brain region, the higher the similarity with the mouse neuron type). Note that we find a near perfect correspondence between mouse neurons and human brain regions.

### In silico modeling of gene networks in AD-relevant neuronal cell types

AD neurodegeneration is the result of multiple molecular-level changes to the system of interacting genes and pathways within vulnerable neurons. We model this system with cell type-specific functional networks, i.e., maps of functional relationships between proteins in the specific cellular contexts of the different types of neurons. Specifically, a functional relationship represents the common involvement of two proteins, either directly or indirectly, in a biological pathway in the cell type of interest. We recently developed a regularized Bayesian network integration method to construct tissue-specific functional networks^36^. These network-level models are an effective first approximation of the functional landscape of a cell and have been successfully applied to the study of diseases^36-38^. It was, however, previously impossible to apply this method to construct networks at neuron-specific resolution because of limitations in high-quality cell type-specific gene expression annotations in human. Given the strong concordance between our mouse neuron-specific molecular signatures and their corresponding human brain regions, we used the signatures as positive examples to extract cell-specific signal from a large human data compendium including thousands of gene expression, protein-protein interaction, and shared regulatory profile datasets to construct human neuron-type specific functional networks. We have made the resulting seven *in silico* human genome-wide network models, each representing one AD-vulnerable or resistant neuron type in the non-disease state, available both for download and dynamic, query-based exploration at http://alz.princeton.edu.

To identify functional characteristics and differences specific to neuron types vulnerable or resistant to AD, we examined the functional cohesiveness of biological processes (i.e., a measure of network connectivity among genes known to be part of that process) in each corresponding functional network model (Supplementary Table 4). We found that pathways neuroprotective in AD^39-41^ appeared more cohesive in AD resistant neurons than in vulnerable neurons, namely the transforming growth factor beta receptor signaling pathway (in DG) and the canonical Wnt signaling pathway (in DG, S1 and V1). On the other hand, mitochondrial processes like apoptotic mitochondrial changes and mitochondrial fission were cohesive in CA1 and ECII respectively, which is consistent with the saliency of mitochondrial dysfunction at early stages of the disease^42^. Strikingly, we found that the processes with largest functional cohesiveness in vulnerable compared to resistant networks were all related to microtubule organization. This is the first evidence that these tau-regulated processes may intrinsically differ between healthy vulnerable and resistant neurons.

### Identifying AD-associated genes through integration of AD GWAS and the ECII functional network

To identify potential AD-associated genes, we then combined these network models of vulnerable neuron function with unbiased disease signal from human quantitative genetics data. Specifically, we developed an approach, Network-Wide Association Study 2.0 (NetWAS 2.0), that extends our previously described^36^ framework with a probabilistic subsampling method to take into account gene-level confidence from quantitative genetics studies. This machine learning approach leverages genome-wide association studies (GWAS) in conjunction with a functional network specific to the region of interest to identify cell type-specific network patterns that are predictive of a disease of interest, reranking all genes based on disease relevance significantly better than the original GWAS^36^.

We applied NetWAS 2.0 using the network model for the most vulnerable neuron (ECII) to reprioritize genes based on an AD GWAS for Braak stage (NFT pathology-based staging)^43^ (Supplementary Table 5). Notably, MAPT (microtubule-associated protein tau, the gene that encodes tau, the primary component of NFTs) was ranked first among all 23,950 reprioritized genes. MAPT was not even nominally significant in the initial GWAS (initial GWAS tau p-value = 0.269). This illustrates the power of NetWAS 2.0 to extract (through cell type-specific functional networks) important disease-relevant signals that may be hidden in the original GWAS. Overall, while the original GWAS for Braak stage was somewhat enriched for known AD genes (genes from the KEGG AD gene set, Wilcoxon rank sum test, p-value < 0.199), the reprioritized gene ranking was much more significantly enriched for these genes (Wilcoxon rank sum test, p-value < 1.60e-4). We also observed strong enrichment of genes involved in regulation of Aβ accumulation and NFT formation (Wilcoxon rank sum test, p-value < 1.29e-10, p-value < 2.2e-16, respectively; gene sets curated by a curator independent from the analyses, Supplementary Table 6) (Fig. 4a, b). Known AD neuroprotective pathways, like neurotrophin signaling^44^ and Wnt signaling pathway^40,41^ were also predicted to be strongly associated (Supplementary Table 7). Lastly, we highlight the association of neurotransmitter secretion with AD (FDR < 2.57e-24). Dysregulation of this pathway is one of the most prominent effects of Aβ accumulation^45^, and the resulting hippocampal network hyperactivity was suggested to be a crucial contributor to AD pathogenesis^46-48^. As the AD signal in NetWAS 2.0 comes only from unbiased GWAS data (i.e., no prior AD disease knowledge was incorporated), the NetWAS 2.0 results thus provide a data-driven, unbiased prioritization of AD-associated processes out of the many pathways that, over time, have been associated with tau pathology.

**Figure 4:**
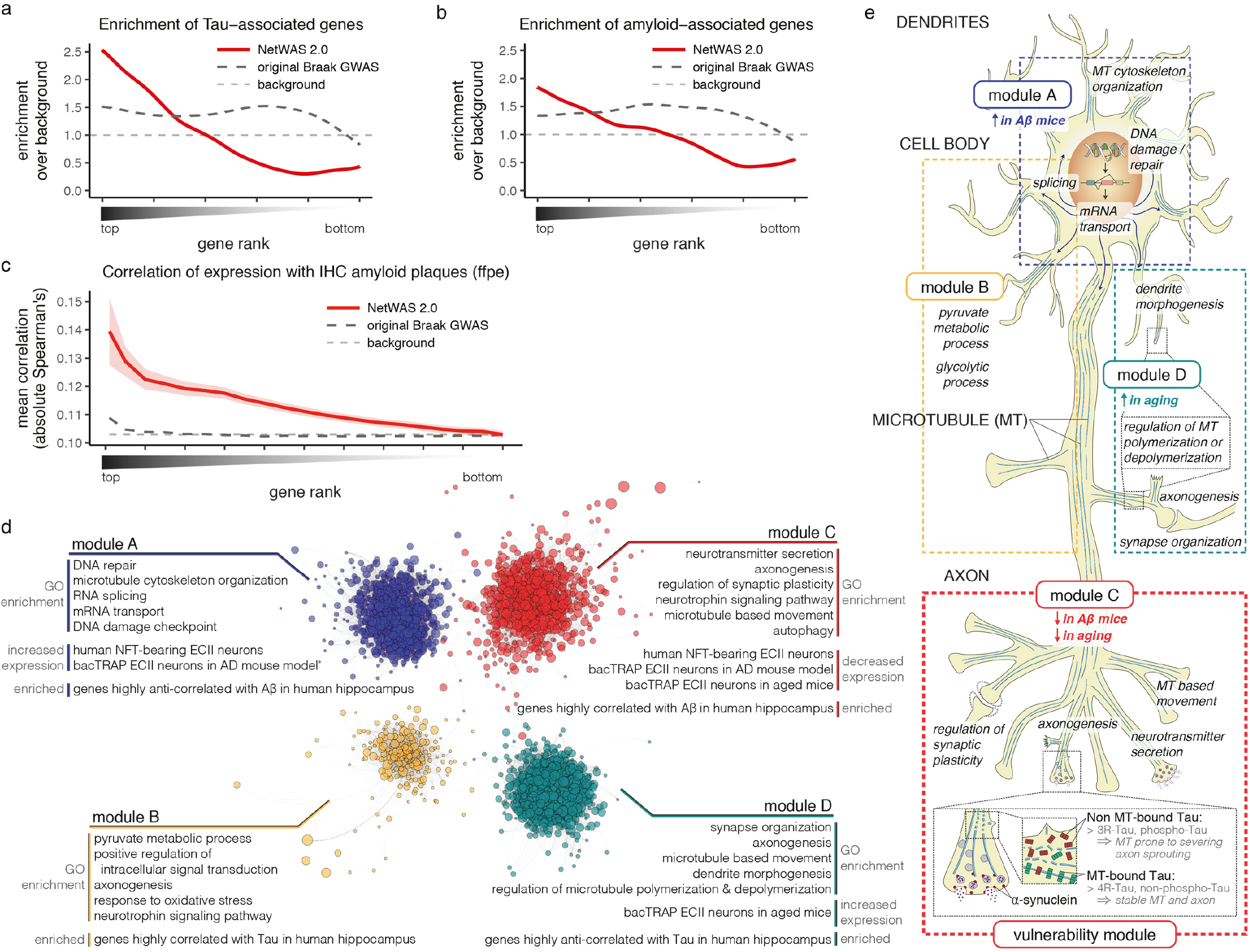
Prediction, validation, and functional analysis of genes associated with AD pathology. **a, b.** Top NetWAS 2.0 gene predictions show significantly higher enrichment of NFT-(a) and Aβ-associated genes (b) (curated by an independent expert, Supplementary Table 6) than the original GWAS used as input to NetWAS 2.0. **c.** Human hippocampal expression levels of top AD-associated gene predictions are highly correlated with amyloid plaque amounts as measured by immunohistochemistry in those subjects^53^. The x-axis represents the proportion of top NetWAS 2.0 genes obtained. The average absolute values of correlations between gene expression level and amyloid plaques across the subset of genes are plotted (NetWAS 2.0 predictions in red with 95% confidence interval; Braak GWAS in black; background genes in grey). **d.** Clustering of the top 10% NetWAS genes using a shared-nearest-neighbor-based community-finding algorithm identifies functional modules corresponding to distinct AD-associated processes. We indicate pathways enriched in each module, as well as the association of each module with aging and AD pathology in both our data (independent from our functional network analysis) and external datasets. Each dot represents a gene (where size inversely correlates with the NetWAS 2.0 ranking, i.e. larger dots represent top ranked genes). Network layout by gephi^96^ of ECII-specific-network posterior probabilities above prior are shown (comembership score ≥ 0.75 based on 1000 subsamples for visual clarity). **e.** Representation of pathways enriched in each module (d) in ECII neurons. Microtubules (MT) are represented in blue. Enrichment for genes modulated by Aβ and aging is indicated for each module. Module A is enriched in neuronal cell body processes, while module C includes many axonal processes. Modules A, B, and D may be generally associated with tau pathology in many types of projection neurons, while module C may capture the surplus of vulnerability from ECII neurons. The module includes both structural and functional axonal remodeling pathways, suggesting that axonal plasticity is key to the degeneration process in AD. Concomitant actions of Aβ and aging on module C genes might perturb crosstalk between axon remodeling processes and eventually impinge on SNCA and MAPT function. Inset: magnified view of an axon terminal. α-synuclein, a regulator of neurotransmitter release, binds to synaptic vesicles (grey circles), to the membrane of the presynaptic active zone, and to MTs. Both forms of tau (3R in red, and 4R in green) are present along MT in the axons, with 4R (as well as non-phosphorylated tau) having higher affinity to MT than 3R (as well as hyperphosphorylated tau). Tau-bound MTs are less stable and more prone to severing, a requirement for axon sprouting and axonal plasticity. We show that PTBP1 regulates both tau isoform usage and α-synuclein levels.

Beyond these well-characterized associations, one of the most significantly enriched pathways in the NetWAS 2.0 results was a microtubule-related process, regulation of microtubule cytoskeleton organization (FDR < 1.38e-27) (Supplementary Table 7). This is consistent with our connectivity analysis, where we discovered that microtubule-regulating pathways were particularly cohesive in vulnerable neurons. Together, these results support a hypothesis that microtubule-regulating pathway genes may cooperate with MAPT for the formation of NFTs in vulnerable neurons. Our data also strongly support a role for mRNA splicing and transport in AD pathogenesis (RNA splicing, FDR = 4.48e-10; RNA transport, FDR = 3.32e-16). RNA binding proteins in these processes have recently emerged as major players in various non-AD neurodegenerative diseases^49^, and recent studies suggested possible involvement in AD (TIA1 protects against tau-mediated degeneration^50^, CELF1 is one of the main GWAS hits^51^, and the activity of ELAVL proteins is altered in AD brains^52^).

### Association of NetWAS 2.0 genes with AD pathology

We next investigated the link between key drivers of the AD pathological cascade (Aβ accumulation and age) and AD-vulnerability-associated genes identified by NetWAS 2.0 analysis. To enable the direct analysis of the ECII-specific effects of Aβ accumulation in AD, we crossed our ECII-bacTRAP mice with an AD mouse model (APP/PS1 mice). These mice overexpress mutant *APP* and *PSEN1* and have increased levels of Aβ in the cortex and hippocampus^53^. We profiled ECII neurons at 6 months of age, when the first plaques are starting to form (Supplementary Table 8). Genes significantly downregulated in these APP/PS1 mice were strongly enriched in our top NetWAS 2.0 gene predictions (Wilcoxon rank sum test, p-value < 9.21e-14). Additionally, genes associated with aging in ECII of wild-type mice (24 month-versus 5-month-old mice) (Supplementary Table 8) were strongly enriched at the top of our ranking (Wilcoxon rank sum test, p-value < 4.19e-13). Our finding that Aβ and aging modulate the expression of genes NetWAS 2.0 predicts to be associated with AD indicates that these genes might connect Aβ accumulation and NFT formation in the age-dependent pathological cascade within vulnerable neurons.

To examine the possible relationship between top NetWAS 2.0 genes and human AD pathology directly, we then used data from two independent human datasets. The Adult Changes in Thought study (ACT)^54^ provides paired gene expression data and pathology measurements from hippocampus samples of elderly individuals at risk for dementia. For each gene, we calculated the correlation between expression level and amount of amyloid plaques. We found that expression of our top gene predictions was significantly more correlated with amyloid plaque amount than either background or genes implicated in the original Braak stage-GWAS (bootstrap p-value < 0.0001, Fig. 4c). Furthermore, our predictions were very significantly enriched in genes differentially downregulated in tangle-bearing ECII neurons of sporadic AD patients measured in a different study (relative to non-tangle-bearing neurons, Wilcoxon rank sum test, p-value < 2.2e-16)^55^. Together, this consensus of results indicates that the top NetWAS 2.0 gene predictions may highlight novel genes that participate in the AD pathological cascade within neurons.

### Identification of AD-associated functional modules

To better understand the processes and pathways through which these genes are associated with NFT formation and AD, we used a shared-nearest-neighbor-based community-finding algorithm^56^ to cluster the genes with top NetWAS 2.0 ranks into functional modules within the ECII network (Fig. 4d, Supplementary Table 9). We identified four modules, each enriched in distinct AD-associated processes, including RNA splicing (module A), metabolism (module B), neurotransmitter release (module C), and neuron differentiation (module D). Interestingly, several pathways were shared across multiple modules, including microtubule organization (A, C, D) and axonogenesis (B, C, D), supporting a central role for these processes in AD pathogenesis (Supplementary Table 9).

We then further characterized these functional modules by examining their relationship to aging as well as Aβ accumulation and NFT formation in vulnerable neurons (Supplementary Table 10). We found that the neurotransmitt er-secretion-related module C genes showed decreased expression in the context of Aβ accumulation in the mouse (our APP/PS1 mouse profiling) and have significantly lower expression in aged wild-type mice. Thus, module C is a good candidate for linking Aβ accumulation with aging in the AD pathological cascade. Furthermore, module C was the only module with ECII-specific signal for tau pathology (i.e., significantly enriched in genes downregulated in NFT-bearing ECII neurons of AD patients, but not strongly correlated with tau in non-ECII regions of the human hippocampal formation (ACT study)). Additionally, only module C demonstrated significantly tighter cohesiveness in ECII versus resistant neurons (Student’s t-test, intersection-union test, p-value < 0.0135). Thus, while modules A, B, and D may represent pathways common to general AD progression in any neuron type, *module C may confer the surplus of susceptibility specific to ECII neurons*. As this vulnerability-specific module represents processes related to both axon structural remodeling and presynaptic excitability, it is tempting to speculate that specific AD vulnerability of ECII neurons may be linked to their lifelong maintenance of a state of high axonal plasticity (Fig. 4e).

### Functional association of α-synuclein, tau, and PTB in ECII neurons

To identify genes in this vulnerability-specific module that underlie ECII susceptibility in relation to early stage AD, we examined the connectivity and centrality of the module members across all seven neuron-specific networks. Intuitively, two genes are tightly connected in a specific neuronal context if they have a high confidence link in the functional network for that neuronal type; this suggests involvement of these genes in shared processes. A highly central gene is one that has many high confidence links across the network, indicating involvement of this gene in a wide array of processes. Within module C, MAPT (tau) was the most centrally connected out of all 668 module C genes, and our analysis pointed to SNCA (α-synuclein) as potentially driving the ECII specificity of this vulnerability-specific module. This is based on the finding that not only are MAPT and SNCA tightly connected to each other in the ECII network, but α-synuclein also has the highest differential network centrality between ECII and the resistant neurons. This suggests that α-synuclein is associated with many more processes in ECII neurons compared to other types of neurons, that tau cooperates with α-synuclein in many of these processes, and that α-synuclein may contribute to NFT formation upon dysregulation of these processes. The novel association between MAPT and SNCA in the context of AD neuronal vulnerability is supported by previous work demonstrating physical as well as functional interaction between these two proteins in other neurodegenerative disorders (reviewed in ^57^). For example, tau and α-synuclein have been previously described to influence each other’s aggregation into pathological lesions in Parkinson’s disease as well as in mice overexpressing these genes^58-61^. However, a role for endogenous α-synuclein in the formation of NFT has not been previously described, although a large proportion of AD patients present α-synuclein pathology^62^.

PTB, a regulator of alternative splicing^63^, was the most highly connected protein to both α-synuclein and tau in the ECII network. Interestingly, PTB was detected in a screen for tau splice factors as one of the regulators of tau exon 10^64^. Regulation of exon 10 is of high relevance for tau pathology, as its inclusion gives rise to four-rather than three-microtubule binding repeat tau (4R- and 3R-tau respectively). An imbalance between 4R- and 3R-tau has been repeatedly shown to give rise to tau pathology in different tauopathies as well as in AD (reviewed in ^65^). Furthermore, NFTs in ECII neurons have been shown to be devoid of 4R-tau, in contrast to other hippocampal neurons that have both tau isoforms^66,67^. Regulation of tau splicing by PTB in ECII neurons could initiate tau pathology specifically in these neurons, potentially contributing to their vulnerability.

## Discussion

Little is known about the molecular basis of selective neuronal vulnerability in AD and the molecular pathways that lead to NFT formation and neurodegeneration. Furthermore, no animal model comprehensively recapitulates every aspect of human AD pathogenesis. Here, we provide an integrative and unbiased framework for the study of this disease that combines advantages of both mouse models and of human data. Our approach 1) models AD vulnerable and resistant human neurons *in silico* with high-quality cell type-specific molecular profiles generated in the non-diseased mouse and a compendium of publicly available human data, 2) leverages human quantitative genetics to identify AD-relevant genes and pathways within these *in silico* models, and 3) experimentally tests in the mouse the effect of age and Aβ, a major AD endophenotype, on the predicted AD genes, elucidating the pathological cascade of AD. Our approach is general and applicable to any complex disease with selective cell vulnerability where relevant human GWAS data are available. For neurodegenerative diseases with a complex multicellular pathogenesis, the approach also allows for the identification of cell type-specific pathological pathways.

Using this approach, we identify molecular mechanisms underlying neuronal vulnerability in AD. In addition to significantly predicting many of the gene candidates previously associated with AD, we also outline novel pathways linking Aβ and tau pathology. Specifically, our unbiased, data-driven analyses place microtubule dynamics at the center of AD pathogenesis. We find that this process is both closely associated with NFT formation and one of the most salient characteristics of the most vulnerable neuronal subtype. As key regulators of neuronal architecture and intraneuronal trafficking, microtubules are the endpoint of many neuronal functional processes. Thus, it is important to determine which specific pathways lead to dysregulation of microtubule dynamics in the context of AD. While a conclusive answer to this question requires further study, our analyses of ECII-vulnerability highlight two potential candidate processes: axonogenesis (which includes tau) and synaptic vesicle release (which includes α-synuclein). Both have been previously linked to microtubule remodeling^68,69^ and are connected to microtubule genes within the vulnerability-specific module. Such interactions could be more prominent in ECII neurons (known to display considerable axon arborization^70^) than in other cell types – which could confer exceptional axonal plasticity to ECII neurons, but could also be responsible for ECII vulnerability.

## Materials and Methods

### Animal models

All experiments were approved by the Rockefeller University Institutional Animal Care and Use Committee (RU-IACUC protocols #07057, 10053, 13645-H), and were performed in accordance with the guidelines described in the US National Institutes of Health Guide for the Care and Use of Laboratory Animals. Mice were housed in rooms on a 12 h dark/light cycle at 22 °C and maintained with rodent diet (Picolab) and water available *ad libitum*. Mice were housed in groups of up to five animals. All bacTRAP mice and APP/PS1 mice (B6.Cg-Tg(APPswe,PSEN1dE9)85Dbo/Mmjax purchased from the Jackson lab) were maintained in a heterozygous state by crossing them with non-transgenic C57Bl/6J mice (also purchased from the Jackson Lab). For cell type-specific profiling in wild-type mice, only male mice were used, and the tissue from two males were pooled. Each type of neuron was profiled at 4-5 months, 12 months, and 24 months. For comparing ECII neurons in wild-type and APP/PS1 mice, both male and female mice were used, and each sample corresponded to the tissue of one mouse.

### bacTRAP transgene construction

In order to construct cell type-specific bacTRAP mice, we searched for drivers specific to each type of neuron. For that purpose, we mined the ABA and GENSAT for genes expressed selectively in the different cell types of interest. We selected the following genes: Rasgrp2 and Sh3bgrl2 (ECII principal neurons), Sstr4 (CA1 pyramidal neurons), Cacng5 (CA2 pyramidal neurons), Gprin3 (CA3 pyramidal neurons), Calca (V1 pyramidal neurons), Cartpt (S1 pyramidal neurons) for enriched expression in the cell type of interest compared to neighboring cell types. Regulatory regions of these genes should drive expression in the corresponding neuron types. We thus used these genes to construct corresponding bacTRAP mice according to previously described procedures^71^. Specifically, we obtained the bacterial artificial chromosomes (BACs) where the open reading frame (ORF) for each of these genes is most centrally located, ensuring that both upstream and downstream regulatory sequences are driving the expression of the bacTRAP construct: RP23-307B16 (Sh3bgrl2), RP23-199D5 and RP24-344N1 (Rasgrp2), RP23-126C5 (SSTR4), RP23-329L1 (Cacng5), RP23-181A2 (Calca), RP24-68J22 (Cartpt) (Children’s Hospital and Research Center at Oakland). We modified each BAC to place the eGFP-L10a cDNA under the control of each gene’s regulatory sequences^71^. For each gene, we cloned by PCR a small homology arm corresponding to approximately 500bp of sequence upstream of the ORF, stopping 5 bp before the ORF (sequences of the small homology arms in supplementary table 11), in the S296 shuttle vector (a pLD53.SC2 plasmid containing the cDNA for eGFP-L10a). For each BAC, we transformed the BAC and the S296 vector containing the corresponding small homology arm into recA-expressing bacteria. We monitored the proper integration of eGFP-L10a at the beginning of each ORF using southern blot. We prepared a purified BAC stock using Cesium Chloride gradient, and linearized the BAC with PI-SceI. The Rockefeller University Transgenic Services performed pronuclear injection of the linearized BACs on a C57Bl/6J (Jackson Lab) background. F1 and F2 of the different founder lines were then tested for proper expression pattern. One of the founder lines with the Sstr4 BAC (Sstr4#19 line) presented ectopic expression in granule cells from the dentate gyrus and no expression in CA1 neurons. We thus used Sstr4#19 for granule cell profiling. We used another founder line (Sstr4#7) for CA1 neurons. We also separately obtained Cck- and Gprin3-bacTRAP mice, which were previously described^26,72^.

### Cell type-specific molecular profiling

To isolate cell type-specific mRNA, bacTRAP mice from the different transgenic lines were decapitated after slight CO_2_ intoxication, and brains were promptly taken out. For each transgenic line, we dissected the minimal area where transgene expression is restricted to the cell type of interest (for ECII bacTRAP lines, we made a coronal cut around -3.3 mm antero-posterior (AP); for Sh3bgrl2-bacTRAP, we then scooped the hippocampus off the tissue caudal to the cut, discarded it, and kept the tissue located ventral to the rhinal fissure; for Rasgrp2-bacTRAP, we took all the tissue caudal to the -3.3 mm AP cut, and ventral to a horizontal cut around -3mm dorso-ventral (DV); for SSTR4#7- and SSTR4# 19-bacTRAP lines, we used all the hippocampus; for the CCK-, CACNG5- and Gprin3-bacTRAP lines, we used all the hippocampus rostral from a coronal cut around -3.3mm AP; for CALCA-bacTRAP, we made a sagittal cut around +3.6 mm medio-lateral (ML) on each side, a coronal cut around -3 mm AP and we extracted the cortex respectively dorsal and caudal to these cuts, while cutting out the mEC; for CARTPT-bacTRAP we made coronal cuts around 1.75 mm AP, -0.25 mm AP, and -2.25 mm AP and for each slice, we dissected out the part of the cortex that contains the somatosensory cortex).

We then performed bacTRAP purification following the previously described procedure^28^ except for two differences. First the volume of lysis buffer used for tissue homogenization depends on the size of each particular brain region. The buffer volumes for each bacTRAP line are shown in Supplementary Table 11. Second, we used RNeasy Plus Micro Kit (Qiagen) to purify RNA after bacTRAP, and RNA was thus detached from beads using the RLT Plus buffer supplemented with 1% β-mercaptoethanol (MP biomedicals). RNA integrity was evaluated with a bio-analyzer RNA 6000 pico chip (Agilent) and RNA quantified by fluorescence detection with Quant-It Ribogreen RNA reagent (ThermoFisher). All samples included in the study had RNA Integrity Numbers above 7. Five ng of RNA were then used for reverse-transcription with Ovation RNAseq v2 kit (NuGEN). cDNAs were cleaned up using a QIAquick PCR purification kit (Qiagen). Doublestranded cDNAs were quantified by fluorescence detection using Quant-IT Picogreen dsDNA reagent (ThermoFisher). cDNAs (200 ng) were sonicated in 120ul volume using a Covaris S2 ultrasonicator (duty cycle, 10%; intensity, 5; cycles/burst, 100; time, 5 minutes) to generate 200bp fragments on average. The fragmented cDNAs were then used to construct sequencing libraries using TruSeq RNA sample prep kit v2 (Illumina). Library concentration was evaluated using bioanalyzer, and libraries were multiplexed. Multiplexes were then sequenced at the Rockefeller University genomics resource center with a HiSeq 2500 sequencer (Illumina).

### Histology

To study the expression pattern of the bacTRAP transgene, bacTRAP mice were transcardially perfused with 4% paraformaldehyde, brains were dissected out, immersion fixed for one hour in 4% paraformaldehyde, frozen in OCT compound (TissueTek), and 40 μm-thick section were cut on a CM3050 S cryostat (Leica). Sections were permeabilized in PBS with 0.1% Fish Gelatin (Sigma), 2% normal goat serum (Jackson ImmunoResearch), and 0.1% triton X-100 and then stained overnight at 4°C in PBS with 0.1% Fish Gelatin and 2% normal goat serum with a chicken anti-GFP antibody (1/300). The primary antibody was detected with an Alex 488-donkey anti-chicken secondary antibody (1/300). After the last wash, sections were mounted with Prolong Gold Medium containing DAPI. Sections were imaged using a Zeiss LSM 510 META laser scanning confocal microscope. Images were minimally processed using Photoshop (Adobe Systems) to enhance brightness and contrast for optimal representation of the data.

### RNA-seq analysis

RNA-sequencing reads were mapped to the mouse genome (Ensembl 75) using STAR (version 2.3.0e, default parameters)^73^, and gene-level counts were quantified using htseq-count (version 0.9.1)^74^. Genes were subjected to an expression detection threshold of 1 count per million reads per gene in more than 3 samples and oligodendrocyte, endothelial, and ependymal cell gene clusters were excluded to focus on the neuronal signal. Differential expression and multidimensional scaling analysis were performed using edgeR (version 3.8.6)^75^.

### Spatial homology analysis

Human brain microarray data were downloaded from the ABA (http://human.brain-map.org/static/download)^76^. Brain regions that were measured in fewer than 3 out of the 6 subjects profiled were excluded from downstream analysis to ensure robustness.

We calculated an ontology-aware spatial homology score between each of our 7 mouse neuron types and each of the 205 human brain regions robustly measured by the ABA, as follows:

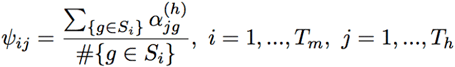, for mouse neuron type *i* and human brain region *j* (thus *T_m_* = 7 and *T_h_* = 205), human gene *g*.

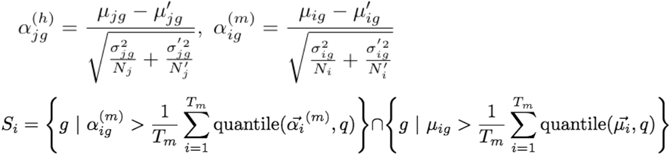

where 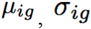 are respectively the mean and standard deviation of expression for the mouse functional ortholog^77^ of gene *g* in mouse neuron type *i* (in log_2_(rpkm)). *N_i_* is the number of samples for neuron type *T_i_* while 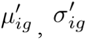, are the mean and standard deviation of expression values for the mouse functional ortholog for unrelated neuron types (e.g., for neuron type hippocampus CA2, 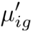 would be the mean expression of all non-hippocampus neuron types). The quantile used was *q* = 0.9. Normalized microarray expression values as processed by the Allen Institute of Brain Science were used to calculate the corresponding scores (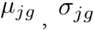, etc.) for gene *g* in human. Intuitively, 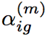 is a normalized enrichment score for the mouse functional ortholog of gene g in tissue *T_i_*, of the mouse. *S_i_* is the set of genes that are both highly expressed (high 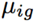) and highly specific (high 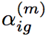) to tissue *T_i_*, thus providing a strong molecular signature for that tissue. This signature is combined with the enrichment scores from human 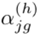 to produce a final spatial homology score.

### Construction of functional networks

We then used these cell type-specific molecular signatures to construct a cell type-specific gold standard (see Gold standard section below), which we then used to integrate a human genome-scale data compendium (see Human data compendium section below) to construct cell type-specific functional networks based on our tissue-specific regularized Bayesian integration method^36^ (see Data integration section below).

### Gold standard

The cell type-specific gold standard was constructed by combining a functional interaction standard and cell type specific signatures. The functional interaction gold standard was constructed based on either the presence or absence of gene co-annotations to expert-selected biological process terms from the Gene Ontology (GO) based on whether the term could be experimentally verifiable through targeted molecular experiments. For each of these 337 selected GO terms, we obtained all experimentally derived gene annotations (i.e., annotations with GO evidence codes: EXP, IDA, IPI, IMP, IGI, IEP). After gene propagation in the GO hierarchy, gene pairs co-annotated to any of the selected terms were considered positive examples, whereas gene pairs lacking co-annotation to any term were considered negative examples, except in cases where the two genes were (i) separately annotated to highly overlapping GO terms (hypergeometric p-value < 0.05) or (ii) coannotated to higher-level GO terms that may still indicate the possible presence of a functional relationship.

We then combined our expanded cell type-specific molecular gene signature sets (q = 0.75) with this functional interaction standard by defining the four classes of edges (C1, C2, C3, and C4) as described in Greene et al^36^, with the adjustment of allowing genes annotated to nervous system tissues to be considered for the C2 negative example class (to emphasize cell type-specificity in relation to other general nervous system genes, rather than excluding them based on the hierarchical tissue ontology as in Greene et al.).

### Human data compendium

We downloaded and processed 31,157 human interaction measurements and brain expression-based profiles from over 24,000 publications, as well as experimentally defined transcription factor binding motifs, chemical and genetic perturbation data, and microRNA target profiles.

Physical interaction data were downloaded from BioGRID (version 3.2.118)^78^, IntAct (Nov 2014)^79^, MINT (2013-03-26)^80^, and MIPS (Nov 2014)^81^. Interaction edges from BioGRID were discretized into five bins (0-4), depending on the number of experiments supporting the interaction. For all other interaction databases, edges were discretized based on the presence or absence of an interaction.

A total of 6,907 expression profiles from 268 human brain expression datasets were downloaded from the Gene Expression Omnibus (GEO)^82^. Duplicate samples were collapsed, and genes with values missing in over 30% of the samples were removed. All other missing values were imputed^83^. Normalized Fisher’s z-transformed expression scores were calculated per pair of genes and discretized into the corresponding bin: (-∞, -1.5), [-1.5, -0.5), [-0.5, 0.5), [0.5, 1.5), [1.5, 2.5), [2.5, 3.5), [3.5, 4.5), [4.5, ∞).

Experimentally defined transcription factor binding motifs were downloaded from JASPAR^84^, and the 1 kb upstream region of each gene was scanned for presence of binding motifs using FIMO^85^ from the MEME software suite^86^. For each pair of genes, the Fisher z-transformed Pearson correlation of binding profiles was calculated and discretized into one of the corresponding bins: (-∞, -1.5), [-1.5, -0.5), [-0.5, 0.5), [0.5, 1.5), [1.5, 2.5), [2.5, 3.5), [3.5, 4.5), [4.5, ∞).

Chemical and genetic perturbation and microRNA target profiles were downloaded from the Molecular Signatures Database (MSigDB, c2:CGP and c3:MIR gene sets, respectively)^87^. For each pair of genes, similarity based on the weighted mean of number of shared profiles (weighted by the specificity of the profile (1/len(genes)) was calculated and discretized into the corresponding bin: (-∞, -1.5), [-1.5, -0.5), [–0.5, 0.5), [0.5, 1.5), [1.5, 2.5), [2.5, 3.5), [3.5, 4.5), [4.5, ∞).

### Data integration

We applied our tissue-specific regularized Bayesian integration method^36^ for each of the 7 neuron types to train a naïve Bayesian classifier by comparing against the positive and negative examples from the cell type-specific gold standard. For each cell type, we constructed a binary class node representing the indicator function for whether a pair of genes have a cell type-specific functional relationship, conditioned on additional nodes representing each of the datasets in the data compendium. Each model was then applied to all pairs of genes in the data compendium to estimate the probability of tissue-specific functional interactions. All code for data integration is available in our open-source Sleipnir library for functional genomics^88^.

### Network connectivity analysis

We calculated a z-score for cohesiveness of various biological process GO terms in each of the neuron-specific networks: 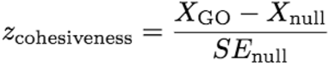 where *X^GO^* is the mean posterior probability of all gene pairs within a particular GO term, and *X_null_*, *SE_null_* are respectively the mean and standard error of the null distribution (based on gene sets randomly sampled within all genes with a GO annotation, with equivalent size to the GO term in question).

### NetWAS 2.0 on AD GWAS

Here, using an AD GWAS for Braak stage (NFT pathology-based staging)^43^ as gold standard and the ECII-specific functional network neighborhoods as features, we applied NetWAS 2.0 with *n*=10,000 to rank each of the 23,950 genes for potential association to AD.

We trained support vector machine classifiers^89^ using (i) nominally significant (p-value < 0.01) GWAS genes as positive examples, (ii) randomly sampled non-significant genes with probability proportional to their GWAS p-value as negatives, and (iii) the network neighborhoods of genes as features. Thus, genes with lower p-values (i.e., more significant) would have a lower chance of being chosen as a negative example than genes with higher p-values. Gene-level p-values were obtained using the versatile gene-based association study 2 (VEGAS2, version:16:09:002) software^90^.

To ensure robustness, we independently sampled *n* such sets of negatives and trained *n* support vector machines. After applying each of the support vector machines to re-rank genes, we aggregated the *n* rankings into a final NetWAS 2.0 gene ranking. Intuitively, the key advance of the NetWAS 2.0 method is that it leverages the GWAS p-values as opposed to treating all non-significant genes as having equal probability of being negative examples as in the original NetWAS method^36^.

### Establishment of the expert-curated gene set

To establish amyloid and NFT gene sets, we recruited a laboratory member, specialized in AD, completely independent from our study, who was unaware of any of the NetWAS 2.0 results. We asked the curator to search for genes involved in tau phosphorylation, aggregation, cleavage, folding, localization, clearance (for the NFT set), and in Aβ production, clearance, aggregation (for the amyloid set). The searches were done with PubMed, and included publications released between January 2000 and April 2017.

### Analysis of NetWAS 2.0 predictions

#### Comparison against the Adult Changes in Thought study

We downloaded paired RNA-seq trascriptomes and neuropathological quantifications from the Adult Changes in Thought (ACT) study (http://aging.brain-map.org/download/index)^54^. We then calculated, for every gene, the Fisher’s z-transformed absolute Spearman’s correlation between its expression in the hippocampus, and IHC amyloid plaque load (ffpe) across all samples.

To aggregate the scores without selecting an arbitrary cutoff, we calculated an amyloid plaque association score for each percentile cutoff averaging the transformed correlation scores for the top x% of NetWAS 2.0 genes (with x=1%, 5%, 10%, 15%, …,100%). We compared these scores against the counterparts calculated based on ranking by the p-values in the Braak GWAS study. For the background distribution, we sampled an equivalent number of genes 1000 times per percentile cutoff.

To calculate bootstrapped 95% confidence intervals for the NetWAS 2.0 amyloid plaque association scores, we subsampled genes with replacement within each percentile cutoff.

### Analysis of Dunckley et al. ECII expression dataset

We downloaded microarray expression profiles measuring tangle-bearing and control LCM ECII neurons ^55^. Data normalization and differential expression analysis were performed using limma (version 3.22.7)^91^. Genes with Benjamini-Hochberg multiple hypothesis test-corrected FDR ≤ 0.05 were considered significantly differentially expressed.

### Identification of functional modules

To identify functional modules represented in our top NetWAS 2.0 genes, we created an ECII subnetwork using the top 10% (i.e., top 2,395) of NetWAS 2.0 ranked genes. Then, we used an approach based on shared *k*-nearest-neighbors (SKNN) and the Louvain community-finding algorithm^56^ to cluster the network into distinct modules. This approach alleviates the effect of high-degree genes and accentuates local network structure by connecting genes that are likely to be functionally clustered together in the ECII network. We calculated the ECII SKNN network by using the number of shared top *k*-nearest neighbors between genes as edge weights and taking the subnetwork defined by the top 5% of edge weights as the subnetwork for downstream analysis. The clustering presented here was calculated with *k*=50, but we confirmed that the clustering was robust for *k* between 10 and 100. Enrichment of Gene Ontology biological process terms and of other experiment-derived gene sets of interest in each module were calculated using one-sided Fisher’s exact tests, with Benjamini-Hochberg multiple hypothesis test correction to calculate FDR.

### Gene connectivity analysis

For each gene *g* in each cell type-specific functional network, we calculated a z-score for gene connectivity, a measure of how central a gene is in the network:
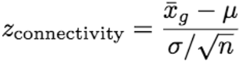 where 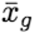 is the average posterior probability of edges incident on gene *g*. μ, σ, and *n* are respectively the mean, standard deviation, and number of all edges in the network.

This is an equation line. Type the equation in the equation editor field, then put the number of the equation in the brackets at right. The equation line is a one-row table, it allows you to both center the equation and have a right-justified reference, as found in most journals.

## End Matter

### Author Contributions and Notes

J.-P.R., V.Y., O.T., and P.G. conceived and designed the research with inputs from M.H. and P.H. J.-P.R. generated mice with help from E.F.S., N.H. and L.B., J.-P.R., Z.P., S.K. and C.A. performed bacTRAP experiments. V.Y. and O.G.T conceived the computational analyses. V.Y. performed all bacTRAP data analyses, generated and analyzed functional networks, reprioritized genes, and re-analyzed publicly available datasets. V.Y. and J.-P.R. analyzed results from the computational analyses. M.F. curated amyloid and NFT lists. J.-P.R., V.Y., O.G.T. and P.G. wrote the manuscript with inputs from M.F. and A.B.C.

## Acknowledgments

We thank Ana Milosevic, Revathy Chottekalapanda, Heike Rebholz, Markus Riessland, Benjamin Kolisnyk, Ruth Dannenfelser, Rachel Sealfon, Chandra Theesfeld, Ran Zhang, for their critical reading of the manuscript, Elisabeth Griggs for assistance with graphic design, transgenic services at Rockefeller University, Bioimaging Resource Center, Genomics Resource Center.

## References

1. Saxena, S. & Caroni, P. Selective neuronal vulnerability in neurodegenerative diseases: from stressor thresholds to degeneration. Neuron 71, 35–48 (2011).

2. Arnold, S. E., Hyman, B. T., Flory, J., Damasio, A. R. & Van Hoesen, G. W. The topographical and neuroanatomical distribution of neurofibrillary tangles and neuritic plaques in the cerebral cortex of patients with Alzheimer’s disease. Cereb. Cortex 1, 103–116 (1991).

3. Fukutani, Y. et al. Neuronal loss and neurofibrillary degeneration in the hippocampal cortex in late-onset sporadic Alzheimer’s disease. Psychiatry Clin. Neurosci. 54, 523–529 (2000).

4. Gómez-Isla, T. et al. Profound loss of layer II entorhinal cortex neurons occurs in very mild Alzheimer’s disease. J. Neurosci. 16, 4491–4500 (1996).

5. Hyman, B. T., Van Hoesen, G. W., Damasio, A. R. & Barnes, C. L. Alzheimer’s disease: cell-specific pathology isolates the hippocampal formation. Science 225, 1168–1170 (1984).

6. West, M. J., Coleman, P. D., Flood, D. G. & Troncoso, J. C. Differences in the pattern of hippocampal neuronal loss in normal ageing and Alzheimer’s disease. Lancet 344, 769–772 (1994).

7. Hof, P. R. & Morrison, J. H. Quantitative analysis of a vulnerable subset of pyramidal neurons in Alzheimer’s disease: II. Primary and secondary visual cortex. J. Comp. Neurol. 301, 55–64 (1990).

8. Morrison, J. H. & Hof, P. R. Life and death of neurons in the aging brain. Science 278, 412–419 (1997).

9. Thal, D. R., Rüb, U., Orantes, M. & Braak, H. Phases of A beta-deposition in the human brain and its relevance for the development of AD. Neurology 58, 1791–1800 (2002).

10. Sepulcre, J. et al. Hierarchical Organization of Tau and Amyloid Deposits in the Cerebral Cortex. JAMA Neurol 74, 813–820 (2017).

11. Braak, H. & Braak, E. Neuropathological stageing of Alzheimer-related changes. Acta Neuropathol. 82, 239–259 (1991).

12. Lewis, D. A., Campbell, M. J., Terry, R. D. & Morrison, J. H. Laminar and regional distributions of neurofibrillary tangles and neuritic plaques in Alzheimer’s disease: a quantitative study of visual and auditory cortices. J. Neurosci. 7, 1799–1808 (1987).

13. Bussière, T. et al. Progressive degeneration of nonphosphorylated neurofilament protein-enriched pyramidal neurons predicts cognitive impairment in Alzheimer’s disease: stereologic analysis of prefrontal cortex area 9. J. Comp. Neurol. 463, 281–302 (2003).

14. Brier, M. R. et al. Tau and Aβ imaging, CSF measures, and cognition in Alzheimer’s disease. Sci Transl Med 8, 338ra66–338ra66 (2016).

15. Arriagada, P. V., Growdon, J. H., Hedley-Whyte, E. T. & Hyman, B. T. Neurofibrillary tangles but not senile plaques parallel duration and severity of Alzheimer’s disease. Neurology 42, 631–639 (1992).

16. Giannakopoulos, P. et al. Tangle and neuron numbers, but not amyloid load, predict cognitive status in Alzheimer’s disease. Neurology 60, 1495–1500 (2003).

17. Ayata, P. et al. Epigenetic regulation of brain region-specific microglia clearance activity. Nat Neurosci 21, 1049–1060 (2018).

18. Cembrowski, M. S., Wang, L., Sugino, K., Shields, B. C. & Spruston, N. Hipposeq: a comprehensive RNA-seq database of gene expression in hippocampal principal neurons. Elife 5, e14997 (2016).

19. Small, S. A. et al. Model-guided microarray implicates the retromer complex in Alzheimer’s disease. Ann. Neurol. 58, 909–919 (2005).

20. Hawrylycz, M. et al. Canonical genetic signatures of the adult human brain. Nat Neurosci 18, 1832–1844 (2015).

21. Liang, W. S. et al. Gene expression profiles in anatomically and functionally distinct regions of the normal aged human brain. Physiol. Genomics 28, 311–322 (2007).

22. Miller, J. A., Oldham, M. C. & Geschwind, D. H. A systems level analysis of transcriptional changes in Alzheimer’s disease and normal aging. J. Neurosci. 28, 1410–1420 (2008).

23. Zhang, B. et al. Integrated systems approach identifies genetic nodes and networks in late-onset Alzheimer’s disease. Cell 153, 707–720 (2013).

24. Mostafavi, S. et al. A molecular network of the aging human brain provides insights into the pathology and cognitive decline of Alzheimer’s disease. Nat Neurosci 21, 811–819 (2018).

25. Heiman, M. et al. A translational profiling approach for the molecular characterization of CNS cell types. Cell 135, 738–748 (2008).

26. Doyle, J. P. et al. Application of a translational profiling approach for the comparative analysis of CNS cell types. Cell 135, 749–762 (2008).

27. Brichta, L. et al. Identification of neurodegenerative factors using translatome-regulatory network analysis. Nat Neurosci 18, 1325–1333 (2015).

28. Heiman, M., Kulicke, R., Fenster, R. J., Greengard, P. & Heintz, N. Cell type-specific mRNA purification by translating ribosome affinity purification (TRAP). Nat Protoc 9, 1282–1291 (2014).

29. Sanz, E. et al. Cell-type-specific isolation of ribosome-associated mRNA from complex tissues. Proc. Natl. Acad. Sci. U.S.A. 106, 13939–13944 (2009).

30. Clarke, L. E. et al. Normal aging induces A1-like astrocyte reactivity. Proc. Natl. Acad. Sci. U.S.A. 115, E1896–E1905 (2018).

31. Sun, S. et al. Translational profiling identifies a cascade of damage initiated in motor neurons and spreading to glia in mutant SOD1-mediated ALS. Proc. Natl. Acad. Sci. U.S.A. 112, E6993–7002 (2015).

32. Zeng, H. et al. Large-scale cellular-resolution gene profiling in human neocortex reveals species-specific molecular signatures. Cell 149, 483–496 (2012).

33. Carlyle, B. C. et al. A multiregional proteomic survey of the postnatal human brain. Nat Neurosci 20, 1787–1795 (2017).

34. Strand, A. D. et al. Conservation of regional gene expression in mouse and human brain. PLoS Genet 3, e59 (2007).

35. Walker, L. C. & Jucker, M. The Exceptional Vulnerability of Humans to Alzheimer’s Disease. Trends Mol Med 23, 534–545 (2017).

36. Greene, C. S. et al. Understanding multicellular function and disease with human tissue-specific networks. Nat Genet 47, 569–576 (2015).

37. Krishnan, A. et al. Genome-wide prediction and functional characterization of the genetic basis of autism spectrum disorder. Nat Neurosci 19, 1454–1462 (2016).

38. Song, A. et al. Network-based analysis of genetic variants associated with hippocampal volume in Alzheimer’s disease: a study of ADNI cohorts. BioData Min 9, 3 (2016).

39. Caraci, F. et al. TGF-beta 1 protects against Abeta-neurotoxicity via the phosphatidylinositol-3-kinase pathway. Neurobiol. Dis. 30, 234–242 (2008).

40. Tesseur, I. et al. Deficiency in neuronal TGF-beta signaling promotes neurodegeneration and Alzheimer’s pathology. J. Clin. Invest. 116, 3060–3069 (2006).

41. Liu, C.-C. et al. Deficiency in LRP6-mediated Wnt signaling contributes to synaptic abnormalities and amyloid pathology in Alzheimer’s disease. Neuron 84, 63–77 (2014).

42. Du, H. et al. Early deficits in synaptic mitochondria in an Alzheimer’s disease mouse model. Proc. Natl. Acad. Sci. U.S.A. 107, 18670–18675 (2010).

43. Beecham, G. W. et al. Genome-wide association metaanalysis of neuropathologic features of Alzheimer’s disease and related dementias. PLoS Genet 10, e1004606 (2014).

44. Nagahara, A. H. et al. Neuroprotective effects of brain-derived neurotrophic factor in rodent and primate models of Alzheimer’s disease. Nat. Med. 15, 331–337 (2009).

45. Abramov, E. et al. Amyloid-beta as a positive endogenous regulator of release probability at hippocampal synapses. Nat Neurosci 12, 1567–1576 (2009).

46. Bakker, A. et al. Reduction of hippocampal hyperactivity improves cognition in amnestic mild cognitive impairment. Neuron 74, 467–474 (2012).

47. Vossel, K. A. et al. Seizures and epileptiform activity in the early stages of Alzheimer disease. JAMA Neurol 70, 1158–1166 (2013).

48. Palop, J. J. & Mucke, L. Amyloid-beta-induced neuronal dysfunction in Alzheimer’s disease: from synapses toward neural networks. Nat Neurosci 13, 812–818 (2010).

49. Ramaswami, M., Taylor, J. P. & Parker, R. Altered ribostasis: RNA-protein granules in degenerative disorders. Cell 154, 727–736 (2013).

50. Apicco, D. J. et al. Reducing the RNA binding protein TIA1 protects against tau-mediated neurodegeneration in vivo. Nat Neurosci 21, 72–80 (2018).

51. Lambert, J. C. et al. Meta-analysis of 74,046 individuals identifies 11 new susceptibility loci for Alzheimer’s disease. Nat Genet 45, 1452–1458 (2013).

52. Scheckel, C. et al. Regulatory consequences of neuronal ELAV-like protein binding to coding and non-coding RNAs in human brain. Elife 5, 4625 (2016).

53. Borchelt, D. R. et al. Accelerated amyloid deposition in the brains of transgenic mice coexpressing mutant presenilin 1 and amyloid precursor proteins. Neuron 19, 939–945 (1997).

54. Miller, J. A. et al. Neuropathological and transcriptomic characteristics of the aged brain. Elife 6, 383 (2017).

55. Dunckley, T. et al. Gene expression correlates of neurofibrillary tangles in Alzheimer’s disease. Neurobiol. Aging 27, 1359–1371 (2006).

56. Blondel, V. D., Guillaume, J.-L., Lambiotte, R. & Lefebvre, E. Fast unfolding of communities in large networks. Journal of Statistical Mechanics: Theory and Experiment 2008, P10008 (2008).

57. Moussaud, S. et al. Alpha-synuclein and tau: teammates in neurodegeneration? Mol Neurodegener 9, 43 (2014).

58. Khandelwal, P. J. et al. Parkinson-related parkin reduces α-Synuclein phosphorylation in a gene transfer model. Mol Neurodegener 5, 47 (2010).

59. Khandelwal, P. J., Dumanis, S. B., Herman, A. M., Rebeck, G. W. & Moussa, C. E.-H. Wild type and P301L mutant Tau promote neuro-inflammation and α-Synuclein accumulation in lentiviral gene delivery models. Mol. Cell. Neurosci. 49, 44–53 (2012).

60. Giasson, B. I. et al. Initiation and synergistic fibrillization of tau and alpha-synuclein. Science 300, 636–640 (2003).

61. Emmer, K. L., Waxman, E. A., Covy, J. P. & Giasson, B. I. E46K human alpha-synuclein transgenic mice develop Lewy-like and tau pathology associated with age-dependent, detrimental motor impairment. J. Biol. Chem. 286, 35104–35118 (2011).

62. Hamilton, R. L. Lewy bodies in Alzheimer’s disease: a neuropathological review of 145 cases using alpha-synuclein immunohistochemistry. Brain Pathol. 10, 378–384 (2000).

63. Llorian, M. et al. Position-dependent alternative splicing activity revealed by global profiling of alternative splicing events regulated by PTB. Nat. Struct. Mol. Biol. 17, 1114–1123 (2010).

64. Wang, J. et al. Tau exon 10, whose missplicing causes frontotemporal dementia, is regulated by an intricate interplay of cis elements and trans factors. J. Neurochem. 88, 1078–1090 (2004).

65. Liu, F. & Gong, C.-X. Tau exon 10 alternative splicing and tauopathies. Mol Neurodegener 3, 8 (2008).

66. Iseki, E. et al. Immunohistochemical investigation of neurofibrillary tangles and their tau isoforms in brains of limbic neurofibrillary tangle dementia. Neurosci. Lett. 405, 29–33

67. Hara, M., Hirokawa, K., Kamei, S. & Uchihara, T. Isoform transition from four-repeat to three-repeat tau underlies dendrosomatic and regional progression of neurofibrillary pathology. Acta Neuropathol. 125, 565–579 (2013).

68. Bodaleo, F. J. & Gonzalez-Billault, C. The Presynaptic Microtubule Cytoskeleton in Physiological and Pathological Conditions: Lessons from Drosophila Fragile X Syndrome and Hereditary Spastic Paraplegias. Front Mol Neurosci 9, 60 (2016).

69. Lewis, T. L., Courchet, J. & Polleux, F. Cell biology in neuroscience: Cellular and molecular mechanisms underlying axon formation, growth, and branching. J. Cell Biol. 202, 837–848 (2013).

70. Tamamaki, N. & Nojyo, Y. Projection of the entorhinal layer II neurons in the rat as revealed by intracellular pressure-injection of neurobiotin. Hippocampus 3, 471–480 (1993).

71. Gong, S. et al. A gene expression atlas of the central nervous system based on bacterial artificial chromosomes. Nature 425, 917–925 (2003).

72. Gray, J. D. et al. Translational profiling of stress-induced neuroplasticity in the CA3 pyramidal neurons of BDNF Val66Met mice. Mol. Psychiatry 23, 904–913 (2018).

73. Dobin, A. et al. STAR: ultrafast universal RNA-seq aligner. Bioinformatics 29, 15–21 (2013).

74. Anders, S., Pyl, P. T. & Huber, W. HTSeq--a Python framework to work with high-throughput sequencing data. Bioinformatics 31, 166–169 (2015).

75. Robinson, M. D., McCarthy, D. J. & Smyth, G. K. edgeR: a Bioconductor package for differential expression analysis of digital gene expression data. Bioinformatics 26, 139–140 (2010).

76. Hawrylycz, M. J. et al. An anatomically comprehensive atlas of the adult human brain transcriptome. Nature 489, 391–399 (2012).

77. Park, C. Y. et al. Functional knowledge transfer for high-accuracy prediction of under-studied biological processes. PLoS Comput. Biol. 9, e1002957 (2013).

78. Chatr-Aryamontri, A. et al. The BioGRID interaction database: 2013 update. Nucleic Acids Res. 41, D816–23 (2013).

79. Orchard, S. et al. The MIntAct project--IntAct as a common curation platform for 11 molecular interaction databases. Nucleic Acids Res. 42, D358–63 (2014).

80. Licata, L. et al. MINT, the molecular interaction database: 2012 update. Nucleic Acids Res. 40, D857–61 (2012).

81. Mewes, H. W. et al. MIPS: curated databases and comprehensive secondary data resources in 2010. Nucleic Acids Res. 39, D220–4 (2011).

82. Barrett, T. et al. NCBI GEO: archive for functional genomics data sets--update. Nucleic Acids Res. 41, D991–5 (2013).

83. Troyanskaya, O. et al. Missing value estimation methods for DNA microarrays. Bioinformatics 17, 520–525 (2001).

84. Mathelier, A. et al. JASPAR 2014: an extensively expanded and updated open-access database of transcription factor binding profiles. Nucleic Acids Res. 42, D142–7 (2014).

85. Grant, C. E., Bailey, T. L. & Noble, W. S. FIMO: scanning for occurrences of a given motif. Bioinformatics 27, 1017–1018 (2011).

86. Bailey, T. L., Johnson, J., Grant, C. E. & Noble, W. S. The MEME Suite. Nucleic Acids Res. 43, W39–49 (2015).

87. Subramanian, A. et al. Gene set enrichment analysis: a knowledge-based approach for interpreting genome-wide expression profiles. Proc. Natl. Acad. Sci. U.S.A. 102, 15545–15550 (2005).

88. Huttenhower, C., Schroeder, M., Chikina, M. D. & Troyanskaya, O. G. The Sleipnir library for computational functional genomics. Bioinformatics 24, 1559–1561 (2008).

89. Joachims, T. A support vector method for multivariate performance measures. in 377–384 (ACM Press, 2005). doi:10.1145/1102351.1102399

90. Mishra, A. & Macgregor, S. VEGAS2: Software for More Flexible Gene-Based Testing. Twin Res Hum Genet 18, 86–91(2015).

91. Smyth, G. K. Linear models and empirical bayes methods for assessing differential expression in microarray experiments. Stat Appl Genet Mol Biol 3, Article3–25 (2004).

92. Saxena, S. & Caroni, P. Selective neuronal vulnerability in neurodegenerative diseases: from stressor thresholds to degeneration. Neuron 71, 35–48 (2011).

