## Supplementary figures and images for "Selective neuronal vulnerability in Alzheimer’s disease: a network-based analysis"

### Supplementary Figure 1

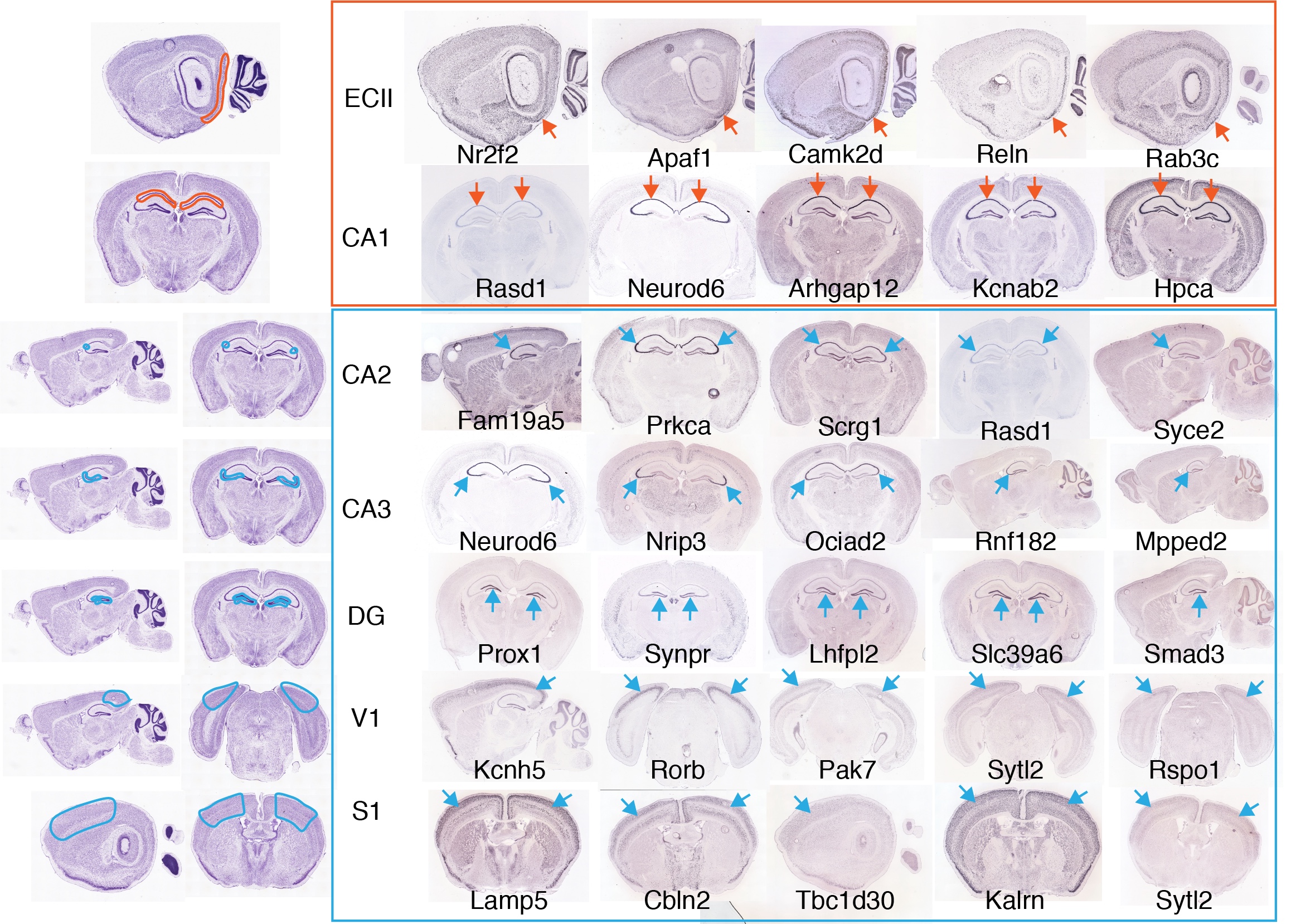
