## Supplementary Note for "Selective neuronal vulnerability in Alzheimer’s disease: a network-based analysis"

**Detail of the bacTRAP mice analyzed:**

Supplementary table 10 lists the bacTRAP lines used for each of the different cell types.

**Mouse genotyping:** All mice were genotyped by PCR on tail clips using the following primers: GFP-forward gacgtaaacggccacaagttcag and GFP-reverse atggtgcgctcctggacgtag for all bacTRAP mice, APP-forward AGGACTGACCACTCGACCAG, APP-reverse CGGGGGTCTAGTTCTGCAT, PSEN1-forward AATAGAGAACGGCAGGAGCA and PSEN1-reverse GCCATGAGGGCACTAATCAT for APP/PS1 mice.

**Comparison of the bacTRAP profiles with Allen Brain Atlas**

In order to cross-validate the bacTRAP profiles, we compared them to Allen Brain Atlas in situ hybridization (ISH) pictures. Based on cell-type specific bacTRAP data, we first calculated an ontology z-score for each gene in each neuron type. For each neuron type, we obtained the 50 genes with the highest ontology z-scores. We then analyzed coronal - whenever possible - or sagittal ISH sections from the Allen Brain Atlas, for each of these genes (50 genes per 7 types of neuron). We scored expression in the seven types of neuron using the Allen Brain Atlas “expression” tool, blind to the identity of the gene, and to the region where it is enriched. Similarly to what Cembrowski *et al.* {Cembrowski:2016df} had done to cross-validate their data, we verified that each gene predicted to be enriched in a given neuron type with our bacTRAP data indeed presented expression in the corresponding neuron type in the Allen Brain Atlas. We found excellent correspondence between our data and the Allen Brain Atlas data, since 98%, 100%, 96%, 100%, 98%, 94% and 87% of the genes predicted to be enriched in ECII, CA1, CA2, CA3, DG, S1 and V1 neurons respectively indeed showed expression in the correct regions in the Allen Brain Atlas (we disregarded genes that do not show expression in any ISH section that are probably expressed below the detection level). In addition, we show detectable signal in our bacTRAP data for some genes with no Allen Brain Atlas ISH signal, like Pkib for example. ISH pictures for the five genes with the highest ontology z-scores for each neuron type, that were available on the Allen Brain Atlas, and that showed expression in some parts of the brain, are shown in supplementary figure 1.

The reference for the Allen Brain Atlas Images shown on Figure 2 and on Supplementary Figure 1 are the following:

| **Figure** | **Gene Symbol** | **Gene name** | **Image series** | **image** |
| --- | --- | --- | --- | --- |
| **Figure 2** | Reln | reelin | 890 | 135 |
|  | Lamp5 | lysosomal-associated membrane protein family, member 5 | 70927827 | 313 |
|  | Whrn | whirlin | 77371813 | 168 |
|  | Wfs1 | Wolfram syndrom 1 homolog (human) | 74881161 | 260 |
|  | Ptpn5 | protein tyrosine phosphatase, non-receptor type 5 | 74743293 | 253 |
|  | Bok | BCL2-related ovarian killer | 71064032 | 252 |
|  | Prox1 | prospero homeobox 3 | 73520980 | 237 |
| **Supp Figure 1** | Nr2f2 | nuclear receptor subfamily 2, group F, member 2 | 308055507 | 9 |
|  | Apaf1 | apoptotic peptidase activating factor 1 | 68745275 | 8 |
|  | Camk2d | calcium/calmodulin-dependent protein kinase II, delta | 68668030 | 5 |
|  | Reln | reelin | 79394359 | 13 |
|  | Rab3c | RAB3C, member RAS oncogene family | 69816745 | 13 |
|  | Rasd1 | RAS, dexamethasone-induced 1 | 2521 | 219 |
|  | Neurod6 | neurogenic differentiation 6 | 698 | 224 |
|  | Arhgap12 | Rho GTPase activating protein 12 | 71836846 | 258 |
|  | Kcnab2 | potassium voltage-gated channel, shaker-related subfamily, beta member 2 | 1754 | 221 |
|  | Hpca | hippocalcin | 72129291 | 248 |
|  | Fam19a5 | family with sequence similarity 19, member A5 | 69059974 | 97 |
|  | Prkca | protein kinase C, alpha | 77869816 | 268 |
|  | Scrg1 | scrapie responsive gene 1 | 71924331 | 268 |
|  | Syce2 | synaptonemal complex central element protein 2 | 70609150 | 74 |
|  | Nrip3 | nuclear receptor interacting protein 3 | 73520999 | 270 |
|  | Ociad2 | OCIA domain containing 2 | 75041527 | 269 |
|  | Rnf182 | ring finger protein 182 | 70719034 | 92 |
|  | Mpped2 | metallophosphoesterase domain containing 2 | 73497744 | 74 |
|  | Prox1 | prospero homeobox 1 | 73520980 | 237 |
|  | Synpr | synaptoporin | 1862 | 229 |
|  | Lhfpl2 | lipoma HMGIC fusion partner-like 2 | 72007934 | 258 |
|  | Slc39a6 | solute carrier family 39 (metal ion transporter), member 6 | 73930852 | 246 |
|  | Smad3 | SMAD family member 3 | 70593360 | 72 |
|  | Kcnh5 | potassium voltage gated channel, subfamily H (eag-related), member 5 | 77620826 | 61 |
|  | Rorb | RAR-related orphan receptor beta | 79556597 | 161 |
|  | Pak7 | p21 protein (Cdc42/Rac)-activated kinase 7 | 75988567 | 165 |
|  | Sytl2 | synaptotagmin-like 2 | 73520979 | 316 |
|  | Rspo1 | R-spondin homolog (Xenopus laevis) | 73636101 | 152 |
|  | Lamp5 | lysosomal-associated membrane protein family, member 5 | 70927827 | 313 |
|  | Cbln2 | cerebellin 2 precursor protein | 70231306 | 300 |
|  | Tbc1d30 | TBC1 domain family member 30 | 72283432 | 1 |
|  | Kalrn | kalirin, RhoGEF kinase | 73930821 | 302 |
